# Applying genomic approaches to identify historic population declines in European forest bats

**DOI:** 10.1101/2023.05.24.542071

**Authors:** Orly Razgour, Cecilia Montauban, Francesca Festa, Daniel Whitby, Javier Juste, Carlos Ibáñez, Hugo Rebelo, Sandra Afonso, Michael Bekaert, Gareth Jones, Carol Williams, Katherine Boughey

## Abstract

1. Anthropogenically-driven environmental changes over the past two centuries have led to severe biodiversity loss, most prominently in the form of loss of populations and individuals. Better tools are needed to assess the magnitude of these wildlife population declines. Anecdotal evidence suggests European bat populations have suffered substantial declines in the past few centuries. However, there is little empirical evidence of these declines that can be used to put more recent population changes into historic context and set appropriate targets for species recovery.
2. This study is a collaboration between academics and conservation practitioners to develop molecular approaches capable of providing quantitative evidence of historic population changes and their drivers that can inform the assessment of conservation status and conservation management. We generated a genomic dataset for the Western barbastelle, *Barbastella barbastellus,* a globally Near Threatened and regionally Vulnerable bat species, including colonies from across the species’ British and Iberian ranges. We used a combination of landscape genetics and approximate Bayesian computation model-based inference of demographic history to identify both evidence of population size changes and possible drivers of these changes.
3. We found that levels of genetic diversity increased and inbreeding decreased with increasing broadleaf woodland cover around the colony location. Genetic connectivity was impeded by artificial lights and facilitated by the combination of rivers and broadleaf woodland cover.
4. The demographic history analysis showed that both the northern and southern British barbastelle populations have declined by 99% over the past 330-548 years. These declines may have been triggered by loss of large oak trees and native woodlands due to shipbuilding during the early colonial period.
5. *Synthesis and applications.* Genomic approaches can be applied to provide a better understanding of the conservation status of threatened species, within historic and contemporary context, and inform their conservation management. This study shows how we can bridge the implementation gap and promote the application of genomics in conservation management through co-designing studies with conservation practitioners and co-developing applied management targets and recommendations.

## Introduction

Anthropogenically-driven environmental changes over the past two centuries have brought on a period of mass global species extinction (WWF, 2018). Yet, the true extent of biodiversity loss is only evident when considering losses of populations and individuals (Ceballos et al., 2017). Wildlife population sizes are estimated to have declined on average by 69% since the 1970s, with highest rates of decline recorded in Latin America and globally in freshwater species (WWF, 2018). The ranges of nearly a third of terrestrial vertebrate species across the globe, including species of low conservation concern, have decreased in the past century, indicating the disappearance of populations (Ceballos et al., 2017). Based on extent of range losses, it has been estimated that many species have already lost more than 10% of their genetic diversity (Exposito-Alonso et al., 2022). By 2050, available habitats for mammals are projected to decline globally by up to 16% due to ongoing climate and land-use changes (Baisero et al., 2020), causing further range contractions and population and genetic diversity losses.

Assessment of wildlife population declines are commonly based on either direct measure of changes in population abundance (WWF, 2018) or indirect indicators, such as proportion of range loss over time (Ceballos et al., 2017). While both approaches offer a good measure of change, they require baseline data, which are not available for all taxa, and in particular all populations. Changes in population size over time (the demographic history of a species) can instead be reconstructed using genomic data obtained from present-day individuals because information on ancestral genomes and the evolutionary forces that shaped them is found within an individual’s genome (Beichman et al., 2018). The advent of high-throughput sequencing technologies over the past 20 years means that genome sequence data are now widely available for nonmodel organisms, including species of conservation concern, and therefore can be used to inform their conservation management (Hohenlohe et al., 2021).

Anecdotal evidence based on range contractions suggests European bat populations have suffered substantial declines in the past few centuries following persecution and extensive land-use changes (Corbet, 1971; Racey & Stebbings, 1972), resulting in habitat loss and reduced prey availability. However, there is little empirical evidence of these declines that can be used to put more recent population changes identified through monitoring efforts in the past 30 years (e.g. Barlow et al., 2015; van der Meij et al., 2015) into historic context and set appropriate targets for species recovery. This lack of strong empirical evidence of historical declines of threatened taxa is a concerning issue in wildlife conservation in Britain (Browning et al., 2021). This study is a collaboration between academics and conservation practitioners aiming to address this issue by developing molecular tools capable of providing quantitative evidence of historic population changes and their drivers that can inform the assessment of conservation status and conservation management.

We focus on the Western barbastelle, *Barbastella barbastellus,* a globally Near Threatened bat species, distributed across Europe, the Caucuses and Morocco (Piraccini, 2016). This bat is classified as Vulnerable in Europe and is considered as a national conservation priority across most European countries. The barbastelle has shown a slight increase in population size in hibernation sites across Europe in the past 20 years (van der Meij et al., 2015). However, globally, their population trend is decreasing, with several reports of local declines (Piraccini, 2016). There are no reliable estimations of population size, even in Britain (Mathews et al., 2018), where there is a national bat monitoring programme (Barlow et al., 2015). The barbastelle is likely to be sensitive to anthropogenic land-use change due to its strong association with mature native woodlands both for roosting and foraging (Russo et al., 2004; Zeale et al., 2012).

We combine landscape genomics tools and model-based inference of demographic history to identify both evidence of population size changes and possible drivers of these changes. We apply our approach to the barbastelle bat, focusing on the British population as a case study, to 1) inform the conservation status of the species, 2) identify landscape and environmental drivers of genetic diversity and connectivity, 3) identify long-term population trends, and 4) show how genomic data can inform the development of conservation management recommendations.

## Materials and methods

### Field sampling

We sampled barbastelle bats (Fig. 1b) during the summers of 2016-2019 in 15 forests across Britain and the Iberian Peninsula, representing parts of the western edge of the species’ range. All forests in Britain were old growth deciduous woodlands, while forests in Spain and Portugal were a combination of deciduous and conifer forests. Bats were caught using mist nets and harp traps with the help of local expert groups. We collected wing biopsy punches from each bat for genomic analysis and stored the samples in RNAlater and -80°C freezers. We included 95 individual bats in the study, 57 from Britain (10 sites) and 38 from Spain and Portugal (five sites; Fig. 1a, Table 1). For population-level analyses (genetic diversity, inbreeding, genetic differentiation and landscape genetics) we only included individuals from sites with five or more individuals and grouped together two sites in Spain with low sample sizes that are within the foraging range of this bat (Zeale et al., 2012) (Table 1). Work was carried out under licences from Natural England, the Home Office, Spanish Administrative regions and The Portuguese Environment Agency, and ethics committee approval of the University of Southampton.

**Figure 1.**
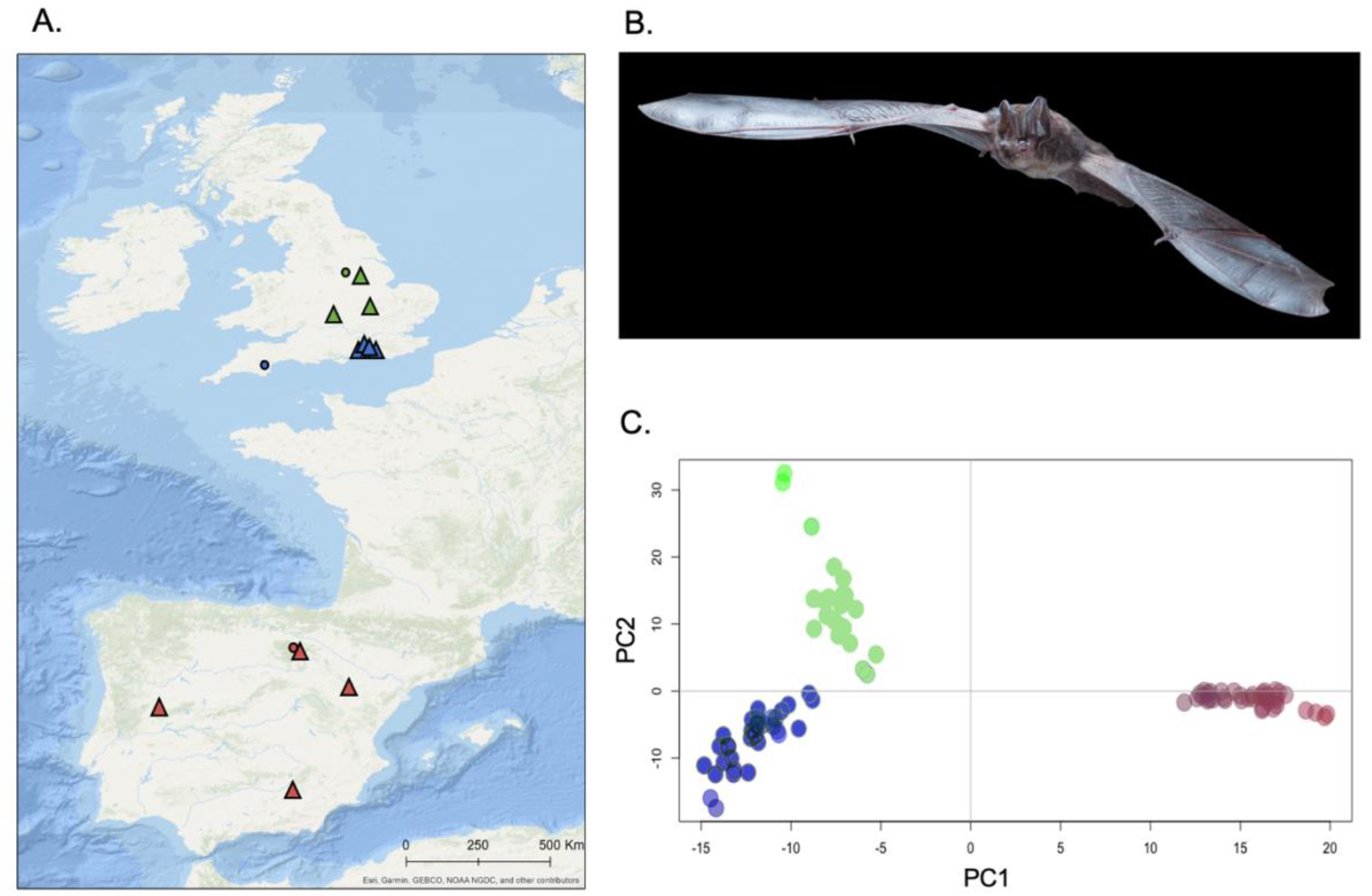
A) Sampling locations for barbastelle colonies marked in triangles and individual locations in circles, colour coded according to their population cluster assignment based on the PCA. B) Photo of the barbastelle in flight (credit: Antton Alberdi). C) Results of the PCA based on genetic distances between individual barbastelle bats. Pink = bats from Spain and Portugal, blue = south of England, green = midlands and north of England.

**Table 1.**
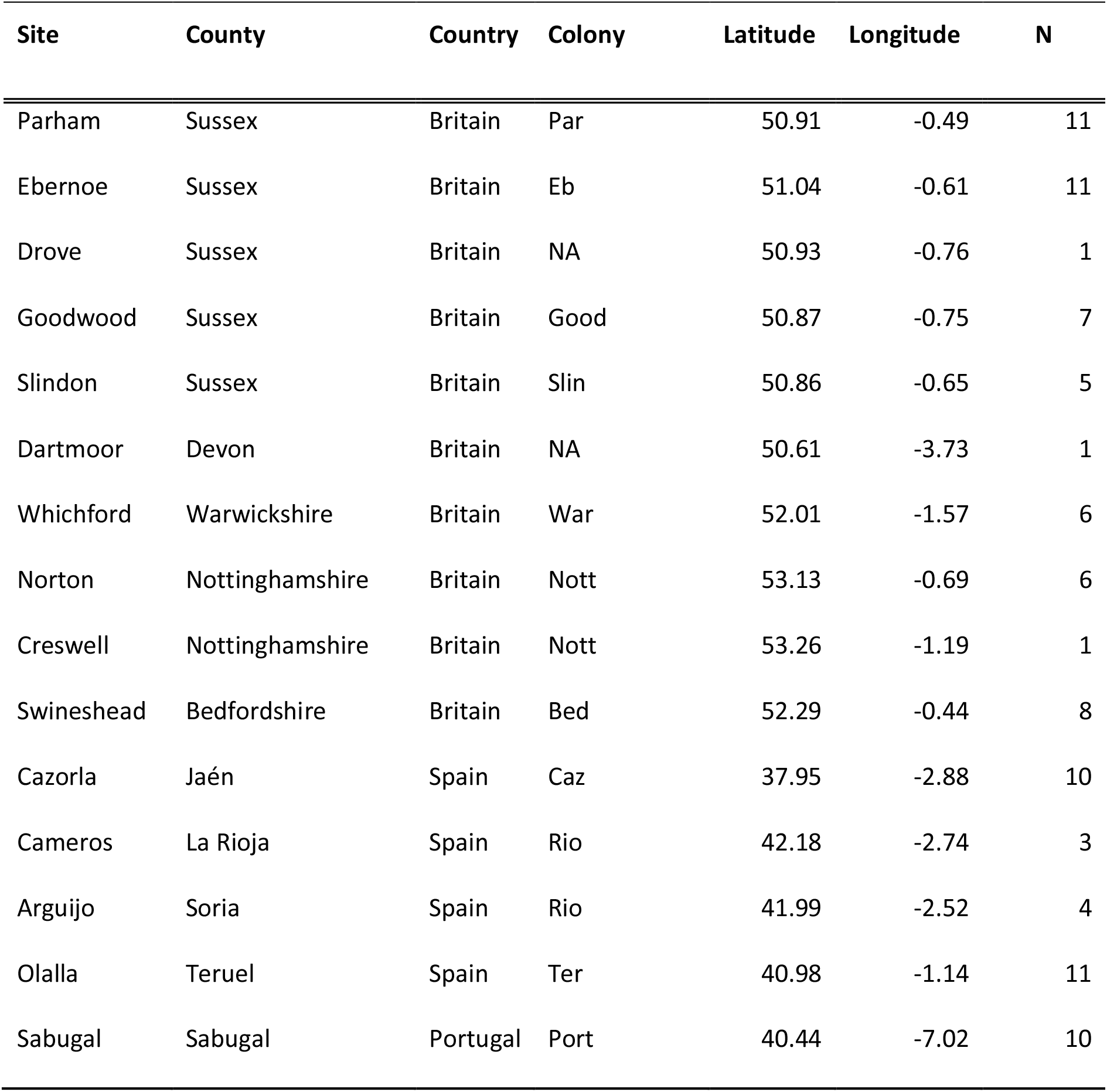
Location and sample sizes (N) of the barbastelle bat sampling sites included in the study, and colony codes.

### Generating the genomic datasets

We used double digest restriction-site-associated DNA sequencing, ddRADseq (Peterson et al., 2012), to generate a genomic dataset containing tens of thousands of anonymous Single Nucleotide Polymorphisms (SNPs) from across the species’ genomes. From the raw sequencing dataset, we generated a high-quality neutral SNP dataset with <5% missing data, including 46,872 SNPs and 95 individual bats. We generated a smaller genomic dataset for the demographic history analysis, including 85 bats and 25,844 SNPs, a sufficient number of SNPs to accurately detect population trends using coalescent-based approaches (Nunziata & Weisrock, 2018) (Supporting Methods for laboratory and bioinformatics procedures).

### Characterising genetic composition

We calculated genetic diversity in Plink v1.9 (Purcell et al., 2007) based on coefficient estimation of observed versus expected individual-level homozygosity (het function) and individual levels of inbreeding based on the correlation between uniting gametes (Fhat3, ibc function). We averaged individual values to generated colony means. We determined population genetic structure based on a PCA on genetic distances, generated using the R package Adegenet (Jombart, 2008). In addition, we used ancestry coefficient plots, generated with the snmf function in the R package LEA (Frichot & François, 2015) to identify the number of population clusters (range 1-10, five replicates) based on cross-entropy values. Following the recommendations in Pearman et al. (2022), we also ran the population structure analysis with a genomic dataset with removed close relatives (IBS>0.499) and retained SNPs out of Hardy-Weinberg equilibrium. We received identical results (Supporting Results). We calculated extent of genetic differentiation between colony pairs using the F_ST_ and Jost’s Dest measures in the R package DiveRsity (Keenan et al., 2013). As values of genetic differentiation based on these measures were highly correlated (R^2^=0.822, P=0.0001), we only include here the results of the F_ST_ analysis.

### Identifying landscape drivers of genetic diversity

We ran linear models and Spearman correlations in R (CRAN) to relate levels of genetic diversity and inbreeding in the bat colonies to landscape variables likely to be important for the species, including percent of broadleaf forest, arable and urban land cover and differences in their percent cover between 1990 and 2019, habitat diversity, and extent of artificial lighting. This analysis was carried out for the full colony dataset (Britain + Iberia), as well as separately for the British colonies. We selected the best supported models based on AICc and adjusted R^2^ values.

### Landscape Genetics analysis

We used the landscape genetics approach to identify barriers to genetic connectivity in the British population. We compared eight candidate models containing resistance values from landscape variables thought to affect genetic connectivity in the barbastelle (Russo et al., 2020) (Table S1-S2; Supporting Methods). Landscape resistance, calculated with Circuitscape v4.0.5 (McRae, 2006), was compared to levels of genetic differentiation (F_ST_) between colonies. As Euclidian geographic distance was highly correlated with genetic distance (F=126.8, R^2^=0.869, P=0.0008), we divided log F_ST_ by log Euclidean distance between colonies to account for the effect of isolation by distance. We used the Maximum Likelihood Population Effect approach (van Strien et al., 2012) and the R packages lme4 (Bates et al., 2015) and usdm (Naimi et al., 2014). To reduce collinearity, candidate models with more than one landscape variable only included variables with VIF<4. The best supported models were identified based on AICc and BIC evidence weights.

### Approximate Bayesian computation inference of demographic history

To identify evidence of historic changes in population size, we reconstructed the demographic history of the British barbastelle populations using the approximate Bayesian computation approach implemented in DIYABC-RF (Collin et al., 2021) (Supporting Methods). Generation length was estimated at 2 years because females commonly reach sexual maturity in their second year (age 1.25 years), but some may become fertile in their first autumn when only a few months old (Russo et al., 2020).

We divided the dataset into three populations: Britain South, Britain North and Spain. We compared four demographic history scenarios (Fig. S1): a null model of no change in population size (S1), recent population decline in both south and north British populations (S2), recent population decline in the south British population with no change in north (S3), and recent population decline in the north British population with no change in the south (S4).

The following priors were used to inform the models: Spanish and British population split dates ranged from pre- to post-LGM (1000-1,000,000 generations ago), split of the British population was set at 10-10,000 to reflect the post-glacial colonisation of Britain, and recent decline dates were set at 10-1000 generations ago. The range of split time between the Iberian and British populations covers more ancient split times to reflect the findings of Rebelo et al. (2012) that Iberia did not contribute to post-glacial colonisation of Europe, including Britain.

## Results

### Population structure

The main population split was between British and Iberian samples. Within Britain, there was a split between north (Bedfordshire, Nottinghamshire and Warwickshire) and south (Sussex and Devon) population clusters. Within the northern cluster, the most differentiated samples were from the most northern colony, Nottinghamshire (Fig. 1c; Fig. S2 for ancestry coefficient plots). Levels of genetic differentiation among colonies were moderate (mean F_ST_=0.048 ±0.02), with highest values (F_ST_>0.07) between British colonies and southern Spain and Portugal, and lowest levels (F_ST_<0.015) between Iberian colonies (Table S3).

### Landscape drivers of genetic diversity

The British colonies, with the exception of Nottinghamshire, had highest levels of heterozygosity and lowest levels of inbreeding (Table S4). In the full dataset, genetic diversity (heterozygosity) was strongly negatively correlated with levels of inbreeding (s=420, P<0.0001, r=-0.909; Fig. S3). Therefore, only results for inbreeding are shown here (Tables S5-S6 for full results). Levels of inbreeding were negatively correlated with percent broadleaf woodlands cover around the colony location (s=410, P=0.0012, r=-0.864) and with present habitat diversity (s=390, P=0.008, r=-0.773; Fig. 2A-B). Among British colonies, levels of inbreeding decreased with increasing broadleaf woodland cover (s=100, P=0.048, r=- 0.786), while genetic diversity tended to increase with percent forest cover around the colony (LM: F=6.34, P=0.053, r=0.748; Fig. 2C-D). Low levels of inbreeding and high levels of heterozygosity were found in areas with >20% broadleaf woodland cover.

**Figure 2.**
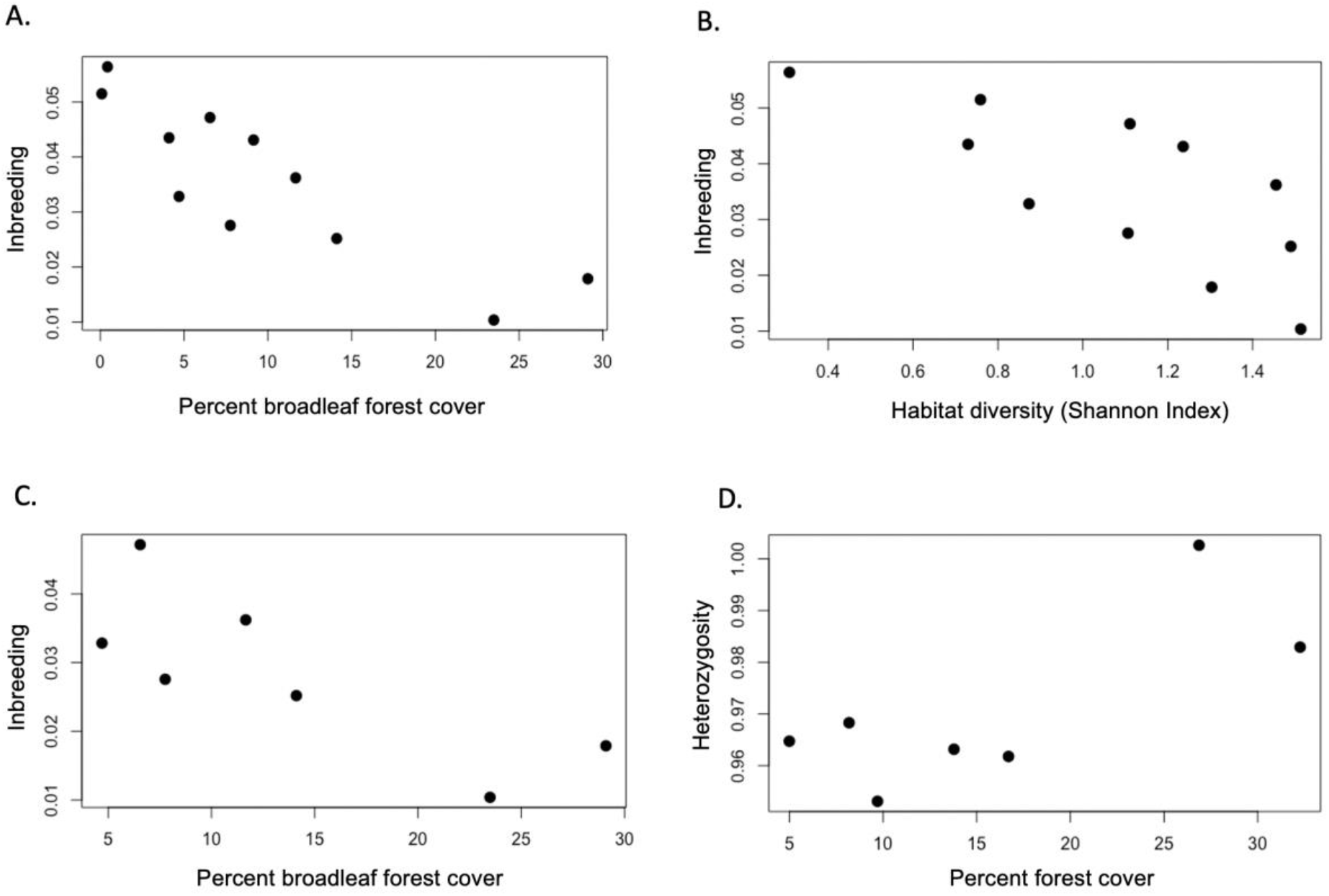
Environmental drivers of genetic diversity in the barbastelle. The relationship between the average levels of inbreeding in British and Iberian colonies and A) percent broadleaf forest cover, and B) current levels of habitat diversity (Shannon Index) in the colony home range, and between average levels of C) inbreeding and D) heterozygosity in the British bat colonies and percent broadleaf forest cover and overall forest cover, respectively.

### Landscape drivers of genetic connectivity

We found evidence for the presence of isolation by distance (IBD) because genetic differentiation among British colonies was highly correlated with geographic distance (F=126.8, R^2^=0.869, P=0.0008). The landscape variables most strongly affecting genetic connectivity between British barbastelle colonies were extent of artificial lights (R^2^=0.432, AICcmi=0.256; connectivity decreased with increasing artificial lights), followed by the combination of rivers and broadleaf woodland cover (R^2^=0.411, AICcmin=0.217; connectivity increased with river and broadleaf cover). The confidence intervals of both models did not overlap zero, supporting their effect on gene flow (Table 2). Movement density maps based on the impacts of artificial lights show limited landscape connectivity around London and other large cities (Fig. 3).

**Figure 3.**
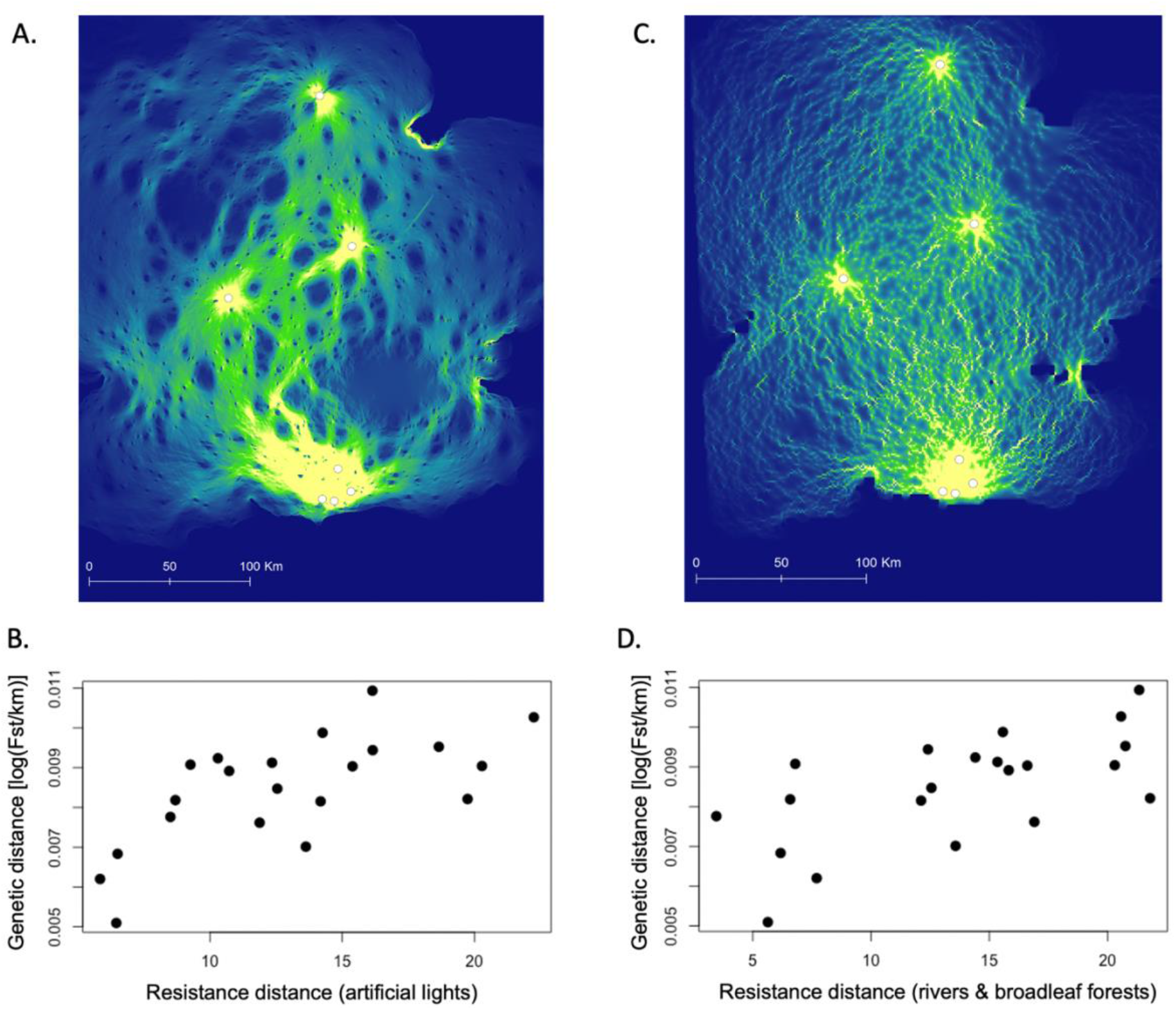
Results of the landscape genetics analysis. Predicted movement density potential between British barbastelle colonies based on the effect of landscape resistance due to A) the intensity of artificial lights at night, and C) rivers and broadleaf woodland cover. Movement density potential ranges from low in dark blue to high in yellow. And the relationship between genetic connectivity (F_ST_ corrected for geographic distance) among British barbastelle colonies and resistance distance due to B) the intensity of artificial lights at night, and D) rivers and broadleaf woodland cover.

**Table 2.** Results of the Maximum Likelihood Population Effect models comparing resistance costs based on different landscape variables to levels of genetic connectivity (F_ST_ corrected for geographic distance) between British barbastelle colonies (AICcmin – Akaike Information Criterion corrected for small sample sizes weights; BICew – Bayesian Information Criterion weights).

| Model | R<sup>2</sup> | AICcmin | BICew | CI 2.5% | CI 97.5% |
| --- | --- | --- | --- | --- | --- |
| Broadleaf woods | 0.363 | 0.169 | 0.165 |  |  |
| Min temp autumn | 0.338 | 0.067 | 0.065 |  |  |
| **River+woods** | **0.411** | **0.217** | **0.212** | 4.47E-05 | 0.0002 |
| River | 0.353 | 0.114 | 0.111 |  |  |
| **Artificial Lights** | **0.432** | **0.256** | **0.250** | 5.70E-05 | 0.0003 |
| Land cover | 0.339 | 0.078 | 0.076 |  |  |
| River+woods & Lights | 0.453 | 0.068 | 0.083 |  |  |
| Woods & Tmin | 0.386 | 0.031 | 0.038 |  |  |

### Model-based inference of demographic history

The observed data fell within the cloud of simulated datasets (Fig. S4). The best supported scenario was a recent population decline in both British populations (scenario S2; posterior probability = 0.573). The demographic history analysis found that the southern and northern Britain populations have declined by 99% around a median of 548 years ago (90% credibility intervals: 126-1066 years ago) and 330 years ago (72-816), respectively. The British population was estimated to split from the Iberian population before the Last Glacial Maximum, a median of 57,682 years ago, while the two British populations split a median of 8,432 years ago, after the post-glacial colonisation of Britain (Table 3; Fig. 4).

**Figure 4.**
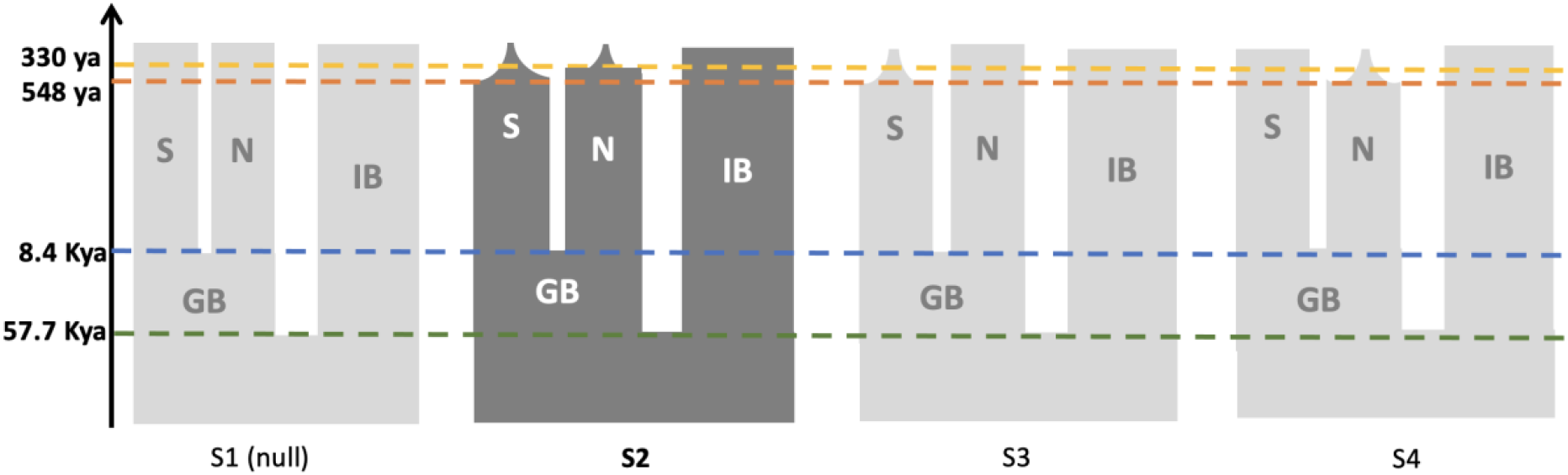
Graphical representation of the four demographic history scenarios compared in the barbastelle demographic history analysis, with the best supported scenario (S2) in dark grey. The median parameter estimations for the best supported scenario are displayed along the y axis (IB – Iberian, GB = British, S = British south and N = British north populations).

**Table 3.**
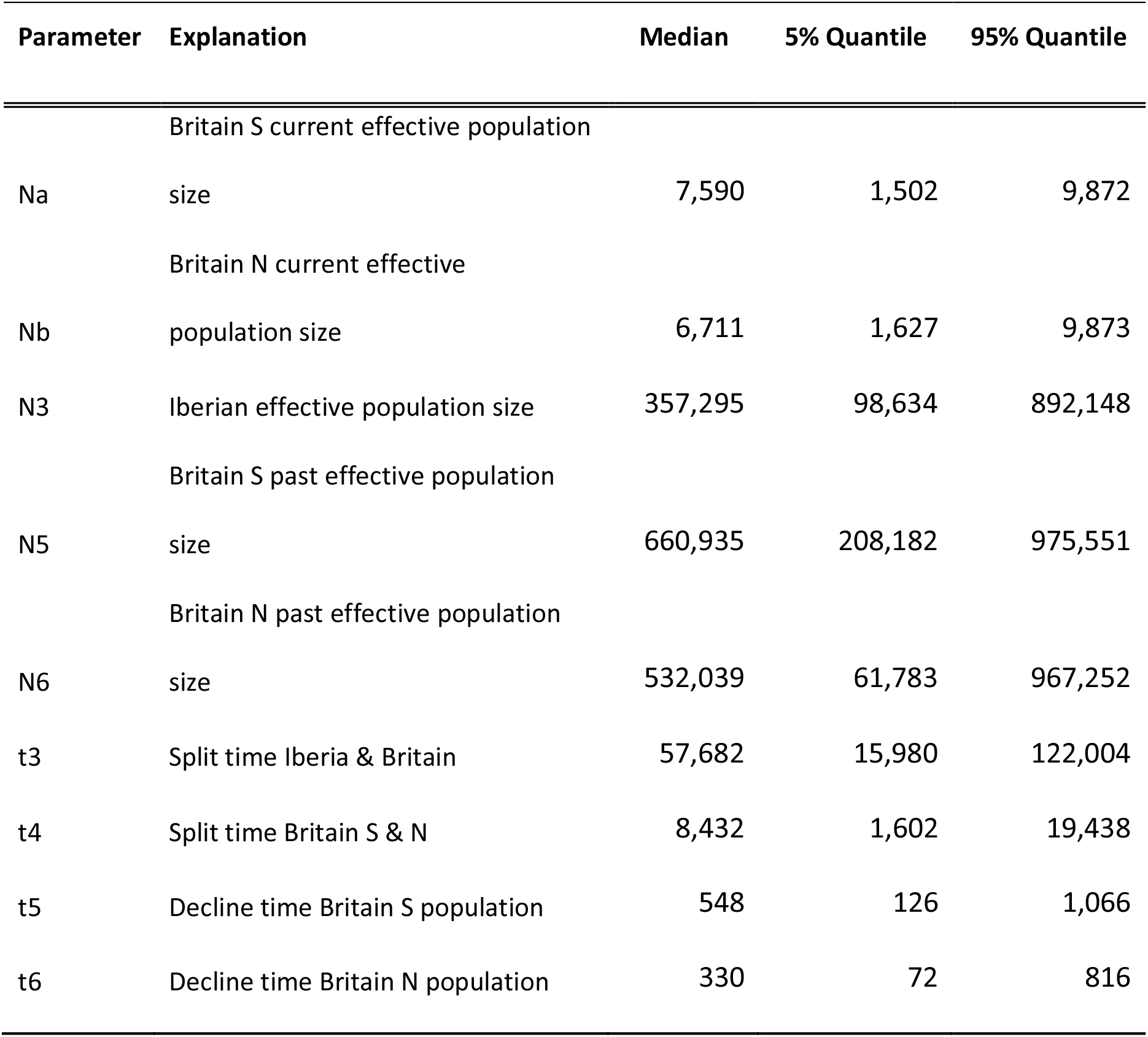
Parameter estimation based on the best supported scenario (S2) in the DIYABC-RF model-based inference analysis (time calculated based on a generation time of 2 years).

## Discussion

We show how genomic approaches can be applied to inform conservation status and identify evidence of population declines in response to anthropogenic land-use changes. Applying landscape genetics tools to the threatened barbastelle bat, we show that genetic composition and population connectivity are affected by broadleaf woodland cover and the extent of artificial lights at night and associated urban expansion. Applying genetically informed model-based inference of demographic history, we identify alarming levels of decline in British barbastelle populations, likely attributes to woodland loss. This decline corresponds to the broader pattern of severe vertebrate population declines across the globe, with 40% of studied European mammals showing ≥80% range contractions in the past century (Ceballos et al., 2017).

### Impacts of anthropogenic land-use change on genetic composition

The barbastelle commonly disperses over distances of up to 75 km, and rarely up to 290 km (Hutterer et al., 2005). Barbastelle bats have relatively long wings and high wing loading values (Norberg & Rayner, 1987), which enables them to fly fast for relatively long distances. Radio-tracked barbastelles in the south of Britain flew to foraging grounds up to 20 km away from the colony roost (Zeale et al., 2012), though typical commuting distances are shorter (Hillen et al., 2009). We found evidence for the presence of isolation by distance among barbastelle colonies in Britain, which support the relatively sedentary nature of this bat (Russo et al., 2020) despite its longer distance dispersal ability.

Barbastelles tend to commute to foraging grounds following natural treelines along watercourses (Greenaway, 2004; Zeale et al., 2012), likely avoiding open areas due to predation risk (Russo et al., 2020). Longer distance dispersal may also be limited due to the strong association of barbastelles with old growth mature broadleaf woodlands both for roosting and foraging (Russo et al., 2004; Zeale et al., 2012), a habitat type that is particularly rare and fragmented in Britain (Reid et al., 2021). In line with these expectations, our landscape genetics analysis identified the combination of rivers and broadleaf woodland cover as a key landscape variable facilitating genetic connectivity between British barbastelle colonies.

Genetic connectivity between colonies is also strongly affected by extent of artificial lights at night. Artificial lights are a key threat to biodiversity that can have negative physiological and demographic consequences (Gaston et al., 2015). Bats are sensitive to artificial lights, but responses vary among species (Voigt et al., 2021). Barbastelles tend to avoid artificially lit areas and their activity is reduced when foraging or drinking sites are artificially illuminated (Russo et al., 2017). Experiments carried out in Britain show that the introduction of artificial lights fragments commuting routes and reduces the activity of other light sensitive bat species (Stone et al., 2012). Hence the continuing expansion of artificial lights in Britain may fragment barbastelle populations and impede genetic connectivity. Artificial lights can be a proxy for other aspects of human disturbance, such as the expansion of urban areas, a habitat type avoided by the barbastelle (Zeale et al., 2012). Lower levels of urbanisation and light pollution across the Iberian Peninsula could explain the higher gene flow rates among Iberian colonies.

Availability of broadleaf woodland cover is a key variable associated with patterns of genetic diversity and inbreeding in barbastelle colonies. Genetic diversity increases and levels of inbreeding decrease with increasing broadleaf woodlands cover around the colony. Summer maternity colonies are mainly found in tree cavities, often in standing dead trees. Barbastelles select relatively tall and wide deciduous trees, usually oak (*Quercus* spp.) or beech (*Fagus sylvatica*), for roosting (Russo et al., 2020). In Britain, barbastelles select scarce standing dead oak trees because they contain more suitable roost cavities (Carr et al., 2018). Maternity colonies on average switch roosts every 3.5 days, and therefore require several large trees that can form an expansive roosting network (Russo et al., 2005). Hence, large mature broadleaf woodlands with veteran and dead trees are needed to support a large enough population to avoid inbreeding and loss of genetic diversity. Many native woodlands have been lost or replaced by plantation forests across Europe, and remaining woodlands are fragmented and degraded (Estreguil et al., 2012). This situation is even more acute in Britain, where forest cover is lower, and the proportion of native and ancient woodlands is particularly low due to historic losses (Reid et al., 2021).

Genetic diversity also increases with increasing habitat diversity across the landscape. Fuentes-Montemayor et al. (2013) found that landscape composition is more important than local woodland characteristics for affecting activity levels of high mobility bat species. Barbastelles forage across the wider landscape, including woodlands, riparian habitats, freshwater wetlands and unimproved grasslands (Zeale et al., 2012). They require a diversity of habitats that can support high densities of moths, their main prey (Goerlitz et al., 2010).

### Broadleaf woodland loss as a driver of population decline

For the first time, using our genomic dataset and model-based inference of demographic history, we were able to provide evidence that confirms suspected historic population declines in the barbastelle bat. Our models show that both the northern and southern British populations have declined by 99% in the past 330 and 548 years, respectively. Despite these severe population declines, contemporary levels of inbreeding are low in British colonies (though slightly higher in the most northern colony, Nottinghamshire), likely reflecting the long-distance movement ability of the species. Moreover, heterozygosity is high relative to Iberian colonies, indicating that despite the severe decline, effective population size is still high enough to prevent substantial losses of genetic diversity (Frankham et al. 2014). This is in line with Díez-del-Molino et al. (2018), who found little concordance between genome-wide diversity and current population sizes in endangered species, likely because the genetic effects of recent population declines are overshadowed by ancient bottlenecks.

The estimated median time of the start of the British barbastelle population declines is set at the mid 15^th^ to late 17^th^ century. This period in European history is referred to as the ‘Age of Discovery’ representing the first wave of European colonialism. Extensive shipbuilding to support colonial explorations required vast quantities of wood (Davey, 2011), and in particular large, ‘outsized’ mature oak trees (Warde, 2006), the same trees favoured by barbastelles for roosting (Russo et al., 2020). Historic documents from that period raise concerns about wood shortage in Britain (Warde, 2006). Although the demand for ship timber likely did not strip the British landscape of wooded resources because it did not involve the full clearing of woodlands, it did result in a shortage of large mature oak trees suitable for shipbuilding, whose rate of regeneration was too slow to meet growing demands (Melby, 2012). Loss of mature native broadleaf woodlands from Britain has continued into the 20^th^ century, with woodland cover dropping from 15% in 1086 to 8% in 1650 and further down to 5.1% in 1924, before increasing to 10% by 2012 (Forestry Commission, 2021). Therefore, the severe decline of the British barbastelle populations may be attributed to loss of maternity colony roosts following the felling of large oak trees for shipbuilding to support colonial explorations and subsequent continued woodland loss.

### Applying genomic approaches to assess historic population declines

Genomic data can improve the accuracy and precision of estimates of population size and demographic history and provide higher power than genetic markers, meaning that the analysis is less limited by sample size (Shafer et al., 2015). However, the type and severity of demographic decline affects the accuracy of the inference, with exponential decline and weak declines being more difficult to identify (Hoban et al., 2014). The severity of the population decline in our case study makes it particularly suitable for genomic inference. Nevertheless, weaker population declines have also been identified using molecular approaches (Razgour et al., 2021). Although more work is needed to assess when approaches to estimate effective population size are reliable, genetic estimates of effective population size can accurately reflect declines in population size (Pierson et al., 2018). Yet, caution should be taken when directly translating effective population size values to actual population sizes for conservation management (Hunter et al., 2018).

Previous applications of genomic approaches to study wildlife population declines have identified more ancient historic declines. For example, Ekblom et al. (2018) found a long-term severe decline in the Scandinavian wolverine population from the onset of the last glaciation, with pronounced decreases in the past 10,000 years. Similarly, Zhao et al., (2012) identified a population bottleneck in the giant panda during the last glacial maximum around 20,000 years ago, with a subsequent decline around 2800 years ago, likely attributed to anthropogenically-driven habitat loss. Our study shows that even recent historic population declines, occurring in the last few hundred years, can be identified using genomic approaches, and therefore can be relevant for informing current conservation management.

### Applying genomic approaches to inform policy and conservation management

The findings of this study are currently being applied in conservation management of the studied species and used to inform policy to support British bats. This study helps place current and future British barbastelle population sizes, trends and indices into context, set appropriate targets for species recovery and develop accurate definitions of Favourable Conservation Status in line with national and international policy obligations.

Genomic approaches can inform our understanding of the historic context for current population estimates and timelines for changes. For most British bat species there is circumstantial evidence for historic loss in the last few centuries based on losses of supporting habitat, insect prey and roosts, coupled with circumstantial evidence. The National Bat Monitoring Programme commenced in 1997 (Barlow et al., 2015), but very few robust datasets exist prior to this. The exception is the greater horseshoe bat, *Rhinolophus ferrumequinum*, where monitoring efforts since the 1940s revealed significant losses and considerable range contraction (Corbet, 1971). It is expected that many other bat species have experienced similar losses due to shared pressures and drivers of change. Lack of a robust historic context has constrained evaluations of the status of bat species and planning for their conservation and recovery into the future. This study shows that genomic approaches can be applied to overcome this constraint in bats and other taxa.

Genomic approaches can further contribute to our ability to assign reasons for decline in population size, reduced genetic diversity and inbreeding. Whilst research in the last 20 years (reviewed in Russo et al. (2020)) has revealed some key landscape, habitat and roosting needs of the barbastelle, these all represent the species survival in an anthropogenically changed landscape. We need a longer term understanding of how species inhabit the landscape and how they have been affected by landscape changes. As we show in this study, genomic data can offer this longer-term perspective.

## Conclusions

Genomic approaches can be applied to inform conservation status and identify evidence of historic population declines in response to anthropogenic land-use change. For the first time we provide evidence that confirms suspected historic population declines in the barbastelle bat. Our models show that both the northern and southern British populations have declined by 99% in the past few hundred years and this decline is likely related to loss of mature broadleaf woodlands due to shipbuilding during the early colonial period. These alarming results place current and future British barbastelle population sizes, trends and indices into context, and are being used to inform recovery targets for this species.

Despite their promises, genomic approaches have not been widely applied in conservation management due to challenges associated with translation into conservation practices and insufficient engagement between academic researchers generating the genomic tools and conservation practitioners (Shafer et al., 2015). This study shows how we can bridge the implementation gap and promote the application of genomics in conservation management through co-designing studies with conservation practitioners and co-developing applied management targets and recommendations. Therefore, this study contributes to better integration of genomic data into national reporting on biodiversity conservation targets and to initiatives to standardise reporting on within species genetic diversity and monitoring of changes (Hoban et al., 2022; O’Brien et al., 2022).

## Authors’ contributions

OR, KB, DW conceived and designed the study. OR, DW, JJ, CI, HR collected the samples. FF, SA, OR did the lab work. MB did the bioinformatics analysis. OR, CM analysed the data. OR, CW, KB led the writing of the manuscript. All authors contributed critically to the drafts and gave final approval for publication.

## Acknowledgements

This project was funded by a Natural Environment Research Council (NERC) Independent Research Fellowship, awarded to OR (NE/M018660) and an award from the Chapman Charitable Trust. We are grateful to the following people for support with fieldwork: Pedro Horta, Helena Raposeira, Francisco Amorim, Diogo Ferreira, Daniel Fernández Alonso, Bob Cornes and Bedfordshire Bat Group, Lois Browne and Warwickshire Bat Group, Matt Cook, Nottinghamshire and Derbyshire Bat Groups, David Rickwood and Andy Carr.

## Conflict of Interest

Razgour is a trustee, and Williams and Boughey are employees of the Bat Conservation Trust (BCT). Whilst this paper details vital findings which will support the conservation work of BCT, there is no source of conflict of interest, financial or otherwise.

## Data Availability

The raw sequencing reads of all libraries will be made available on EBI/ENA. Study locations are presented in Table 1. SNP datasets in vcf format are being uploaded to Dryad.

