## Supplementary Information for "Applying genomic approaches to identify historic population declines in European forest bats"

### Supporting Methods

#### Laboratory procedures

We extracted genomic DNA from the 95 barbastelle bat samples using the Qiagen DNeasy Blood and Tissue extraction kit. DNA was quantified with Qubit® Fluorometer 2.0 and the dsDNA High Sensitivity assay kit (Invitrogen, ThermoFisher Scientific). ddRAD library preparation protocol was based on (Peterson et al., 2012). DNA samples were digested by two high fidelity restriction enzymes at equal ratios: SbfI (CCTGCA|GG recognition site), and MseI (T|TAA recognition site) (New England Biolabs, UK). Libraries were sent for sequencing at the Novagene European Genomic Facility using the Illumina NovaSeq platform with pair-end sequencing.

#### Generating the genomics datasets

We used Plink v1.9 (Purcell et al., 2007) to process the SNP dataset. We filtered out SNPs that had more than 5% missing data (geno 0.05) and minor allele frequencies below 3% (maf 0.03), and SNPs that were out of Hardy-Weinberg equilibrium (hwe 1E-5). The filtered dataset included 46,939 SNPs and 95 individual bats, with a total genotyping rate of 0.992. To generate a neutral genomic dataset, we ran outlier scans in BayeScan (Foll & Gaggiotti, 2008) with 1 million iterations, 50,000 burn-in and 20 pilot runs, to identify loci under directional selection. We removed 67 SNPs (final neutral dataset included 46,872 SNPs). This dataset was used for the population structure, genetic composition and landscape genetics analyses.

We generated smaller genomic datasets for the demographic history analysis that only included SNPs with 99% coverage and minor allele frequencies above 5%. We removed from this dataset the Portugal colony because it was unlikely to be the source of colonisation for the British population. The smaller genomic dataset included 85 bats and 25,844 SNPs.

#### Variables for landscape drivers of genetic diversity analysis

We used the CEH 1990 and 2019 UK land cover map (https://www.ceh.ac.uk/data/ukceh-land-cover-maps) and the ESA CCI 1992 and 2018 land cover map (https://www.esa-landcover-cci.org) to calculate the percent cover of broadleaf woodland, all woodland, urban and arable land covers within the colony sustenance zone of 6 km radius around sampling sites (Collins, 2016). Analysis was carried out in ArcGIS v10.8 (ESRI). We calculated habitat diversity based on Shannon's index and percent difference in woodland cover between 1990 and 2019 (for British colonies) or 1992 and 2018 (for Iberian colonies).

#### Generating resistance surfaces for the Landscape Genetics analysis

We converted landscape variables to resistance surfaces in ArcGIS, assigning different resistance costs based on knowledge of the ecology of this species and its movement behaviour. Resistance costs ranged from one, no resistance to movement, to 100, strong barrier to movement (Table S1). We used Circuitscape v4.0.5 (McRae, 2006) to calculate resistance distance matrices between the British colonies based on the cumulative cost of movement due to landscape resistance. We first used multiple regression on distance matrices in the R package ecodist (Goslee & Urban, 2007) to select the best combination of resistance costs for each landscape variable based on the strength of correlation with genetic distance ($F_{ST}$) between colonies (Table S2).

#### ABC inference of demographic history

We used the approximate Bayesian computation approach implemented in DIYABC-RF (Collin et al., 2021). For each analysis we generated around 25,000 simulations per scenario (100,000 in total). We used random forest analysis to identify the best supported scenario and estimate the value of the different parameters in that scenario. The model choice random forest analysis included 100,000 simulated datasets, 500 trees, 50 summary statistics, three axes of summary statistics LDA linear combination and five noise variables. The parameter estimation analyses included 5054 simulated datasets, 500 trees, minimum node size of 5, 50 summary statistics, 19 axes of summary statistics PLS linear combination, five noise variables and 10,000 out-of-band samples used as test.

### Supporting Results

#### Population structure analysis with the revised dataset

The genomic dataset excluding close relatives and including SNPs out of Hardy-Weinberg equilibrium included 90 individuals and 46,968 SNPs. Similar to the original SNP dataset, the main population split was between British and Iberian samples. Within Britain, there was a split between north and south population clusters, both based on the PCA (Fig. S5) and the snmf ancestry coefficient analysis (Fig. S6).

### Supporting Figures

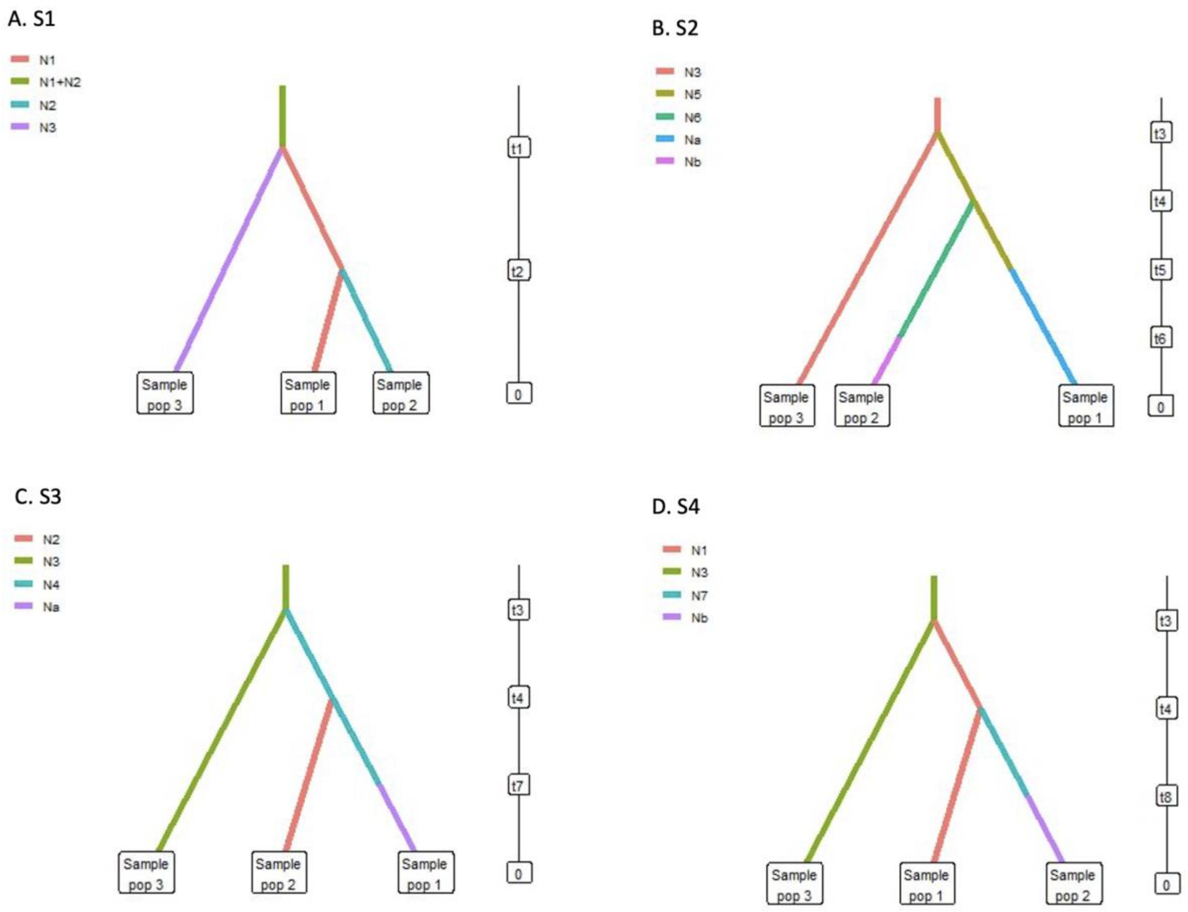

**Fig. S1** – The four demographic history scenarios for the barbastelle bat British population compared in the DIYABC-RF analysis (pop 1 = Britain South, pop 2 = Britain North, pop 3 = Spain).

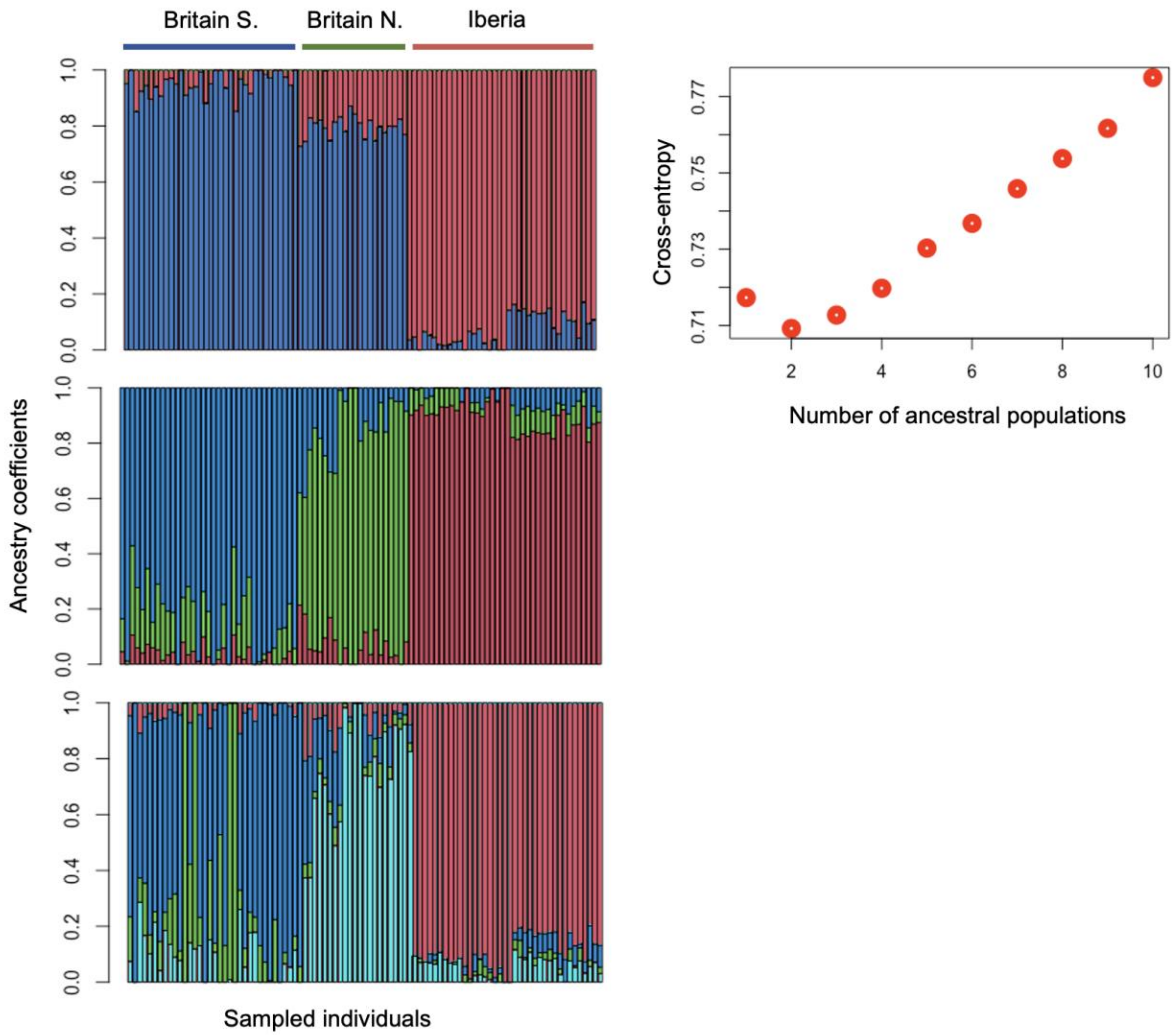

**Fig. S2** – Barbastelle samples divided based on their ancestry coefficients in each population cluster. Each bar represents an individual bat (Britain S. = samples from the south of England, Britain N. = samples from the midlands and north of England, Iberia = samples from Spain and Portugal). Cross-entropy plots showing the likely number of ancestral populations based on lowest cross-entropy values (2 and 3).

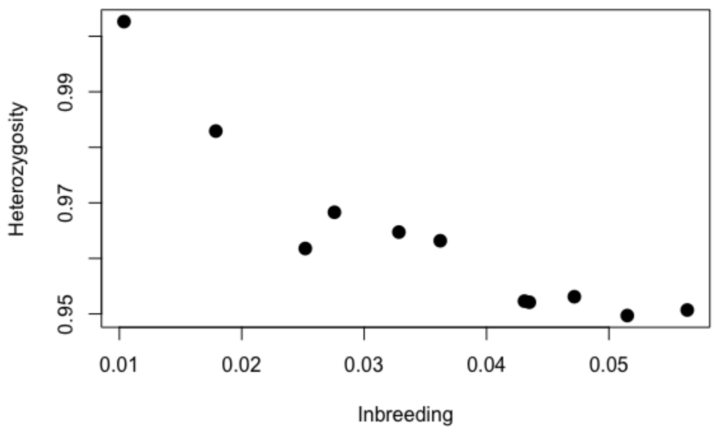

**Figure S3** - The relationship between the average levels of heterozygosity and inbreeding in barbastelle colonies.

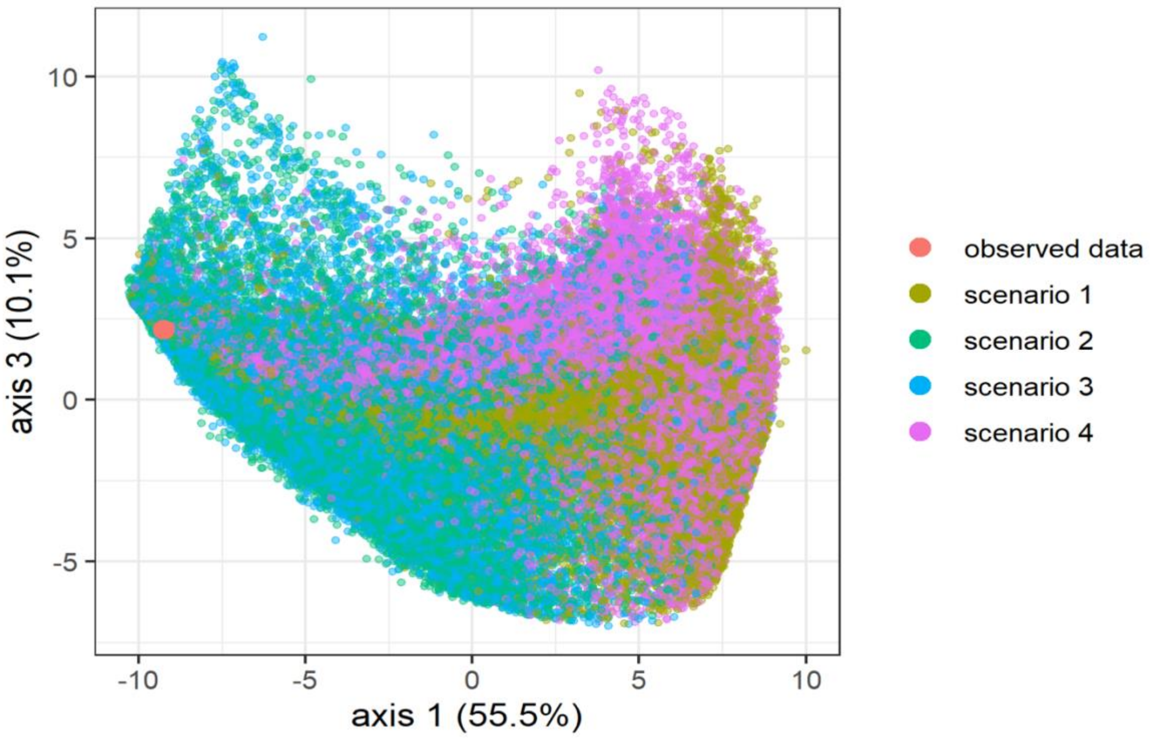

**Figure S4** – PCA showing where the observed data (pink) falls relative to simulated datasets based on the four demographic history scenarios for the British barbastelle populations.

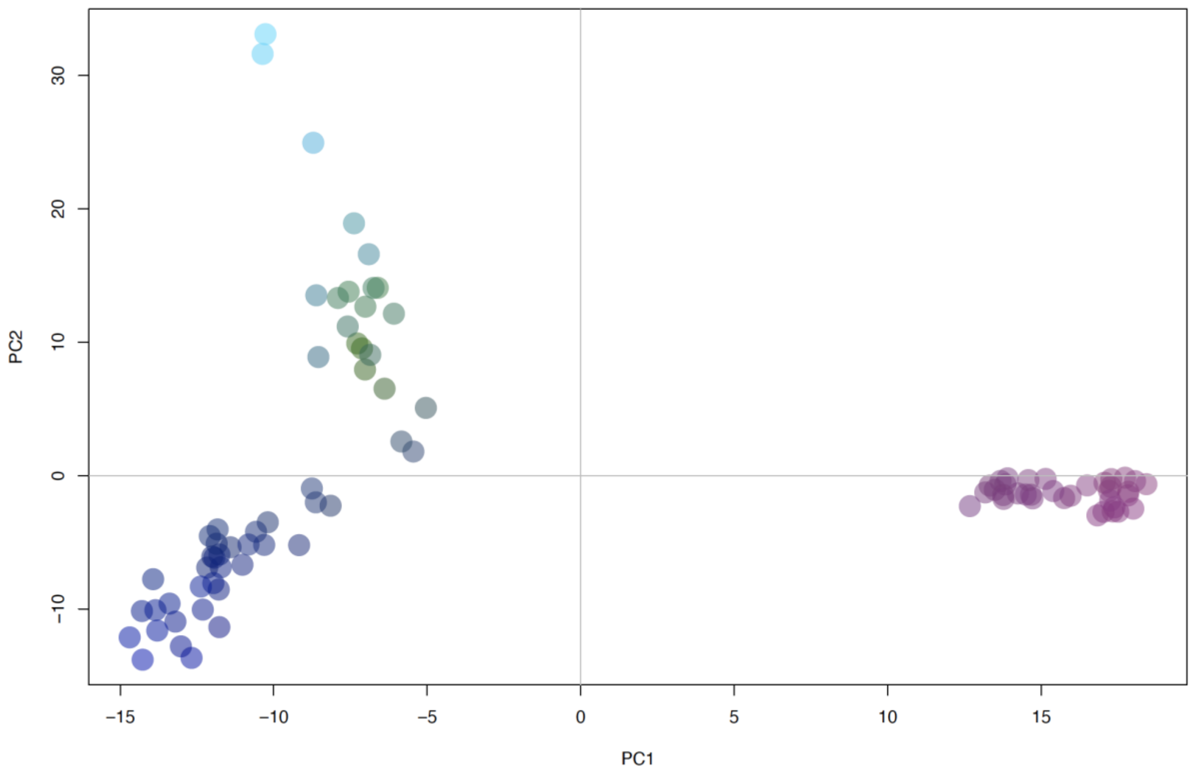

**Figure S5 –** PCA on the genetic distances between individual barbstelle bats based on the revised genomic dataset excluding close relatives and including loci out of Hardy-Weinberg equilibrium. Pink = bats from Spain and Portugal, dark blue = bat from the south of England, light blue-green = bats from the midlands and north of England.

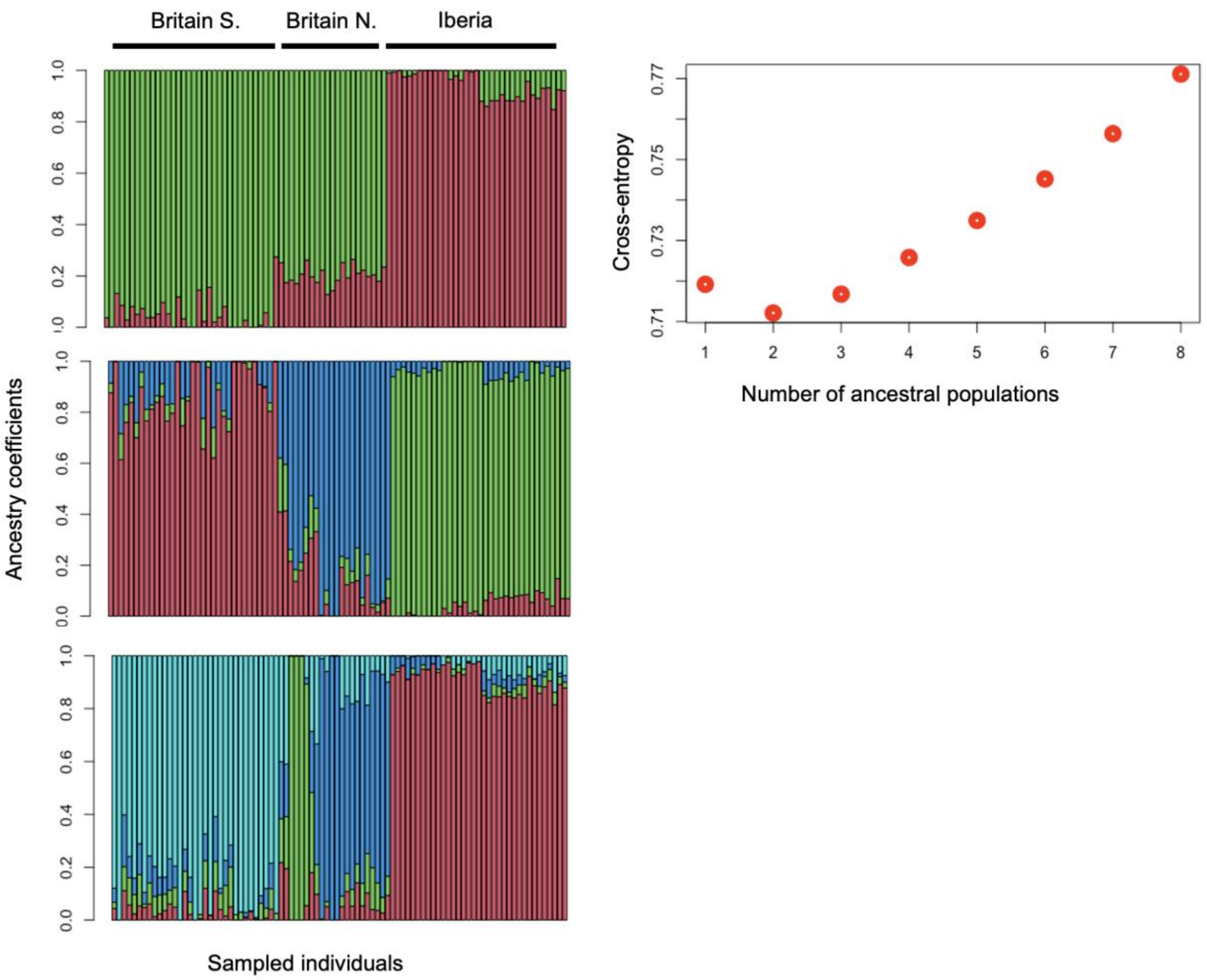

**Figure S6** – Barbastelle samples divided according to their ancestry coefficients in each population cluster, based on the revised genomic dataset excluding close relatives and including loci out of Hardy-Weinberg equilibrium. Each bar represents an individual bat (Britain S. = samples from the south of England, Britain N. = samples from the midlands and north of England, Iberia = samples from Spain and Portugal). Cross-entropy plots showing the likely number of ancestral populations based on lowest cross-entropy values (2 and 3).

### Supporting Tables

**Table S1** – List of variables included in the landscape genetics analysis, their sources, effect on movement with rationale, and assigned resistance costs.

| Variable | Source map | Effect on movement (rationale) | Resistance costs |
|---|---|---|---|
| Land cover 1 | CEH 2019 UK land cover map (www.ceh.ac.uk/ data/ukceh-land-cover-maps) | Strong effect of land cover on gene flow, strongest effect urban. Only main foraging grounds, forests and grasslands, do not form strong barriers. Broadleaf woodland facilitate gene flow more than conifer forest. | 7 categories:<br>broadleaf = 1<br>conifer forest = 10<br>natural grass = 20<br>improved grass = 30<br>arable = 40<br>water, coastal, mountain = 50<br>urban = 100 |
| Land cover 2 | CEH 2019 UK land cover map (www.ceh.ac.uk/ data/ukceh-land-cover-maps) | Strong effect of urban cover on gene flow. Other land cover weaker effect, reflecting the bats flying ability. Broadleaf woodland facilitate gene flow more than conifer forest. | 5 categories:<br>broadleaf woodland = 1<br>conifer forest = 10<br>grass, arable, mountain, water = 20<br>coastal = 50<br>urban = 100 |
| Land cover 3 | CEH 2019 UK land cover map (www.ceh.ac.uk/ data/ukceh-land-cover-maps) | Simplified. Medium effect of land cover, with urban forming the strongest barrier and no difference between forest types | 3 categories:<br>broadleaf, conifer = 1<br>all other land cover = 30<br>urban = 50 |
| Land cover 4 | CEH 2019 UK land cover map (www.ceh.ac.uk/ data/ukceh-land-cover-maps) | Medium effect of land cover, with urban forming the strongest barrier, followed by coastal and mountain. No difference between forest types | 4 categories:<br>broadleaf, conifer = 1<br>grass, arable, water = 20<br>coastal, mountain = 30<br>urban = 50 |
| Land cover 5 | CEH 2019 UK land cover map (www.ceh.ac.uk/ data/ukceh-land-cover-maps) | Simplified. Weak effect of land cover, with urban forming stronger barrier and no difference between forest types | 3 categories:<br>broadleaf, conifer = 1<br>all other land cover = 10<br>urban = 30 |

| | | | |
|---|---|---|---|
| Distance to broadleaf woodland | National Forest Inventory 2018 woodland map (https://data-forestry.opendata.arcgis.com/) | The greater the distance from woodland the less likely the bats are to fly through there | 1-100 (resistance increases with distance from woodlands) |
| Distance to all forest | CEH 2019 UK land cover map (www.ceh.ac.uk/data/ukceh-land-cover-maps) | The greater the distance from woodlands & forests the less likely the bats are to fly through there | 1-100 (resistance increases with distance from woodlands & forests) |
| Percent tree cover | (https://earthenginepartners.appspot.com/science-2013-global-forest/download_v1.2.html) | Resistance to movement decreases with tree cover<br>(tend to follow trees when flying) | 1-100 (resistance increases with decreasing tree cover) |
| River 10 | ESRI (www.arcgis.com/home/item.html?id=273980c20bc74f94ac96c7892ec15aff | Bats tend to commute along rivers, so rivers facilitate movement. Other land cover weak effect on gene flow. | 2 categories:<br>river buffers = 1<br>remaining landscape = 10 |
| River 50 | ESRI (www.arcgis.com/home/item.html?id=273980c20bc74f94ac96c7892ec15aff | Bats tend to commute along rivers, so rivers facilitate movement. Other land cover medium effect on gene flow. | 2 categories:<br>river buffers = 1<br>remaining landscape= 50 |
| Broadleaf woodland & rivers 10 | Merged broadleaf woodland and river maps | Bats roost and forage in woodlands and commute along rivers, therefore these will facilitate gene flow. Weak effect of other land covers. | 2 categories:<br>river buffers, broadleaf = 1<br>remaining landscape = 10 |
| Broadleaf woodland & rivers 50 | Merged broadleaf woodland and river maps | Bats roost and forage in woodlands and commute along rivers, therefore these will facilitate gene flow. Stronger effect of other land covers. | 2 categories:<br>river buffers, broadleaf = 1<br>remaining landscape = 50 |
| Distance to broadleaf woodland & rivers | Merged broadleaf woodland and river maps | Bats roost and forage in woodlands and commute along rivers. | 1-100 (resistance increases with distance from woodlands and rivers) |

| | | | |
|---|---|---|---|
| Night lights | NASA Black Marble (https://blackmarble.gsfc.nasa.gov) | Bats tend to avoid lit areas, therefore resistance increasing with increasing light intensity | 1-100 (resistance increases with increasing light intensity) |
| Human footprint | NASA Last of the wild v2 (https://sedac.ciesin.columbia.edu/data/set/wildareas-v2-human-footprint-geographic/data-download) | Bats tend to avoid areas with high human disturbance, therefore resistance increasing with increasing light intensity | 1-100 (resistance increases with increasing human disturbance) |
| Autumn minimum temperatures | CHELSA Climate, Sep & Oct Tmin (https://chelsa-climate.org/) | Bat disperse and fly to swarming sites for mating during the autumn. Bats are not active at low temperatures. | 1-100 (resistance increases with decreasing temperatures) |

**Table S2** – Results of MRDM tests for associations between genetic ($F_{ST}$) and geographic (Euclidean) distance and to select the best combination of resistance costs for each variable group (hypothesis tested in the landscape genetic analysis) based on their relationship with genetic differentiation ($F_{ST}$) corrected for geographic distance ($F_{ST}$/km) between British barbastelle colonies. Selected variables are highlighted in green.

| Variable group | Variables | $R^2$ | F | P |
|---|---|---|---|---|
| **Distance** | **Fst ~ Euclidean distance** | **0.869** | **126.8** | **0.0008** |
| Land cover | Fst/km ~ Land cover 1 | 0.317 | 8.81 | 0.068 |
| Land cover | Fst/km ~Land cover 2 | 0.234 | 5.79 | 0.102 |
| Land cover | Fst/km ~ Land cover 3 | 0.339 | 9.77 | 0.061 |
| Land cover | Fst/km ~ Land cover 4 | 0.319 | 8.92 | 0.072 |
| Land cover | Fst/km ~ Land cover 5 | 0.301 | 8.19 | 0.082 |
| Forest | Fst/km ~ Distance to broadleaf woodland | 0.363 | 10.84 | 0.035 |
| Forest | Fst/km ~ Distance to all forest | 0.305 | 8.35 | 0.041 |
| Forest | Fst/km ~ Percent tree cover | 0.298 | 8.07 | 0.099 |
| River | Fst/km ~ River 10 | 0.353 | 10.39 | 0.007 |
| River | Fst/km ~ River 50 | 0.262 | 6.75 | 0.010 |
| River & woodlands | Fst/km ~ Broadleaf woodland & rivers 10 | 0.249 | 6.32 | 0.116 |
| River & woodlands | Fst/km ~ Broadleaf woodland & rivers 50 | 0.191 | 4.48 | 0.162 |
| River & woodlands | Fst/km ~ Distance to broadleaf woodland & rivers | 0.411 | 13.26 | 0.01 |
| Anthropogenic | Fst/km ~ Night lights | 0.432 | 14.44 | 0.008 |
| Anthropogenic | Fst/km ~ Human footprint | 0.272 | 7.11 | 0.037 |
| Temperature | Fst/km ~ Autumn minimum temperatures | 0.338 | 9.70 | 0.053 |

**Table S3** – Levels of genetic differentiation, based on $F_{ST}$ values (bottom triangle) and Jost's D values (top triangle), among barbastelle bat colonies in Britain and Iberia (see Table 1 for colony names and locations). Cells are colour-coded from lowest values in green to highest values in red.

| | Par | Eb | Good | Sli | War | Not | Bed | Caz | Por | Rio | Ter |
|---|---|---|---|---|---|---|---|---|---|---|---|
| **Par** | | 0.0000 | 0.0003 | 0.0000 | 0.0007 | 0.0013 | 0.0007 | 0.0026 | 0.0023 | 0.0013 | 0.0020 |
| **Eb** | 0.0118 | | 0.0001 | 0.0000 | 0.0006 | 0.0013 | 0.0007 | 0.0024 | 0.0020 | 0.0011 | 0.0018 |
| **Good** | 0.0267 | 0.0196 | | 0.0000 | 0.0008 | 0.0015 | 0.0011 | 0.0030 | 0.0026 | 0.0015 | 0.0022 |
| **Sli** | 0.0126 | 0.0143 | 0.0158 | | 0.0004 | 0.0008 | 0.0005 | 0.0019 | 0.0016 | 0.0009 | 0.0014 |
| **War** | 0.0385 | 0.0398 | 0.050 | 0.0462 | | 0.0012 | 0.0006 | 0.0022 | 0.0019 | 0.0011 | 0.0016 |
| **Not** | 0.0463 | 0.049 | 0.0585 | 0.0514 | 0.0554 | | 0.0008 | 0.0033 | 0.0031 | 0.0019 | 0.0027 |
| **Bed** | 0.0361 | 0.0387 | 0.0493 | 0.0425 | 0.0419 | 0.0421 | | 0.0023 | 0.0022 | 0.0012 | 0.0019 |
| **Caz** | 0.0592 | 0.059 | 0.0715 | 0.0651 | 0.064 | 0.0734 | 0.0602 | | 0.0000 | 0.0000 | 0.0001 |
| **Por** | 0.0581 | 0.0574 | 0.0693 | 0.0628 | 0.0635 | 0.0746 | 0.0627 | 0.0123 | | 0.0000 | 0.0000 |
| **Rio** | 0.0461 | 0.0458 | 0.0573 | 0.0519 | 0.0517 | 0.0615 | 0.0492 | 0.0105 | 0.0081 | | 0.0000 |
| **Ter** | 0.0506 | 0.0506 | 0.0613 | 0.0562 | 0.0559 | 0.066 | 0.0552 | 0.0154 | 0.0131 | 0.0093 | |

**Table S4** – Levels of genetic diversity in barbastelle colonies in Britain and the Iberian Peninsula, based on mean heterozygosity and inbreeding (see Table 1 for colony names and locations). Cells are colour-coded from lowest levels of heterozygosity and highest levels of inbreeding in red to highest levels of heterozygosity and lowest levels of inbreeding in green.

| Colony | Sample size | Mean Heterozygosity | Mean inbreeding |
|---|---|---|---|
| Par | 11 | 0.979 | 0.021 |
| Eb | 9 | 0.98 | 0.021 |
| Goo | 7 | 0.962 | 0.033 |
| Slin | 5 | 1 | 0.004 |
| War | 6 | 0.978 | 0.020 |
| Not | 7 | 0.936 | 0.065 |
| Bed | 8 | 0.964 | 0.036 |
| Caz | 10 | 0.952 | 0.046 |
| Port | 8 | 0.951 | 0.050 |
| Rio | 7 | 0.954 | 0.043 |
| Ter | 10 | 0.952 | 0.045 |

**Table S5** – Results of the landscape drivers of genetic diversity analysis for heterozygosity and inbreeding in the British colonies, based on Spearman correlations (A) and linear models (B).

| **A. Spearman correlations** | | | |
|---|---|---|---|
| **Models** | **S** | **P** | **r** |
| **Het~IBC** | **100** | **0.048** | **-0.786** |
| **ibc~broadleaf** | **100** | **0.048** | **-0.786** |
| Het~broadleaf | 26 | 0.236 | 0.536 |
| Het~urban | 74 | 0.497 | -0.321 |
| Het~arable | 80 | 0.354 | -0.428 |
| Het~Hab_div2019 | 46 | 0.713 | 0.178 |
| Het~hab_div1990 | 52 | 0.906 | 0.071 |
| Het~diff_broad | 54 | 0.964 | 0.036 |
| Het~diff_arable | 64 | 0.782 | -0.143 |
| Het~diff_urban | 32 | 0.354 | 0.429 |
| ibc~forest | 94 | 0.109 | -0.679 |
| ibc~arable | 28 | 0.267 | 0.500 |
| ibc~urban | 60 | 0.906 | -0.071 |
| ibc~hab_div2020 | 86 | 0.236 | -0.536 |
| ibc~hab_div1990 | 70 | 0.595 | -0.250 |
| ibc~diff_broad | 44 | 0.661 | 0.214 |
| ibc~diff_arable | 36 | 0.444 | 0.357 |
| ibc~diff_urban | 68 | 0.662 | -0.214 |
| ibc~light | 38 | 0.498 | 0.321 |
| **B. Linear models** | | | |
| **Models** | **F** | **P** | **r** |
| **Het~forest** | **6.341** | **0.0533** | **0.747** |
| ibc~light_median | 2 | 0.216 | 0.535 |

**Table S6** – Results of the landscape drivers of genetic diversity analysis for heterozygosity and inbreeding in the full dataset, based on Spearman correlations.

| Models | S | P | r |
|---|---|---|---|
| **Het_Av~IBC_Av** | **420** | **<0.0001** | **-0.909** |
| **IBC_Av ~ perc_broadleaved** | **410** | **0.001** | **-0.864** |
| **LN_IBC ~ Hab_div_2019** | **390** | **0.008** | **-0.773** |
| **Het_Av~perc_broadleaved** | **46** | **0.006** | **0.791** |
| **Het_Av~hab_div2019** | **72** | **0.028** | **0.673** |
| IBC_Av~hab_div1990 | 354 | 0.052 | -0.609 |
| IBC_Av ~ diff_broadleaved | 142 | 0.286 | 0.354 |
| IBC_Av~diff_arable | 310 | 0.214 | -0.409 |
| IBC_Av ~ perc_forest | 140 | 0.273 | 0.363 |
| IBC_Av~perc_arable | 286 | 0.371 | -0.300 |
| IBC_Avg~light | 152 | 0.357 | 0.308 |
| IBC_Avg~light_median | 191 | 0.697 | 0.132 |
| Het_Av~diff_arable | 102 | 0.097 | 0.532 |
| Het_Av~perc_forest | 342 | 0.082 | -0.554 |
| Het_Av~diff_broadleaves | 244 | 0.755 | -0.109 |
| Het_Av~Light | 303 | 0.253 | -0.377 |
| Het_Av~Light_median | 248 | 0.707 | -0.128 |
| Het_Av~perc_arable | 126 | 0.193 | 0.427 |
| Het_Av~hab_div1990 | 116 | 0.146 | 0.473 |
